# Perturbed neurochemical and microstructural organization in a mouse model of prenatal opioid exposure: a multi-modal magnetic resonance study

**DOI:** 10.1101/2023.02.23.529659

**Authors:** Syed Salman Shahid, Gregory G. Grecco, Brady K. Atwood, Yu-Chien Wu

## Abstract

Methadone-based treatment for pregnant women with opioid use disorder is quite prevalent in the clinical environment. A number of clinical and animal model-based studies have reported cognitive deficits in infants prenatally exposed to methadone-based opioid treatments. However, the long-term impact of prenatal opioid exposure (POE) on pathophysiological mechanisms that govern neurodevelopmental impairment is not well understood. Using a translationally relevant mouse model of prenatal methadone exposure (PME), the aim of this study is to investigate the role of cerebral biochemistry and its possible association with regional microstructural organization in PME offspring. To understand these effects, 8- week-old male offspring with PME (n=7) and prenatal saline exposure (PSE) (n=7) were scanned in vivo on 9.4 Tesla small animal scanner. Single voxel proton magnetic resonance spectroscopy (^1^H-MRS) was performed in the right dorsal striatum (RDS) region using a short echo time (TE) Stimulated Echo Acquisition Method (STEAM) sequence. Neurometabolite spectra from the RDS was first corrected for tissue T1 relaxation and then absolute quantification was performed using the unsuppressed water spectra. High-resolution in vivo diffusion MRI (dMRI) for region of interest (ROI) based microstructural quantification was also performed using a multi-shell dMRI sequence. Cerebral microstructure was characterized using diffusion tensor imaging (DTI) and Bingham-neurite orientation dispersion and density imaging (Bingham-NODDI). MRS results in the RDS showed significant decrease in N-acetyl aspartate (NAA), taurine (tau), glutathione (GSH), total creatine (tCr) and glutamate (Glu) concentration levels in PME, compared to PSE group. In the same RDS region, mean orientation dispersion index (ODI) and intracellular volume fraction (VF_IC_) demonstrated positive associations with tCr in PME group. ODI also exhibited significant positive association with Glu levels in PME offspring. Significant reduction in major neurotransmitter metabolites and energy metabolism along with strong association between the neurometabolites and perturbed regional microstructural complexity suggest a possible impaired neuroadaptation trajectory in PME offspring which could be persistent even into late adolescence and early adulthood.

## Introduction

As the opioid epidemic has progressed over the past two decades, opioid use disorder has increased in pregnant women leading to an alarming rise in the number of babies being born with prenatal opioid exposure (POE) and undergoing neonatal opioid withdrawal [1–3]. Prenatal exposure to opioids produces adverse health consequences at birth with substantial neurobehavioral disruptions that persist at least into adolescence if not longer [2–4]. Clinical neuroimaging studies reveal macrostructural and morphometric disturbances in infants with POE, as well as profound alterations in functional connectivity [5]. However, there are substantial differences in the outcomes reported in clinical studies, likely due to the interaction of POE with environmental factors [4]. Animal models of POE are therefore especially valuable to disentangle environmental factors from the effects of the POE itself.

Using prenatal rodent models of opioid exposure, a number of studies have reported neurodevelopmental alterations in the offspring of the exposed animals [6]. However, preclinical neuroimaging studies on opioid exposure are rather limited. To the best of our knowledge, there are only six rodent neuroimaging studies to date which have reported no macrostructural changes [7], alterations in major white matter tracts [8], widespread microstructural changes [9, 10] and changes in cerebral metabolites [11, 12]. There is a knowledge gap regarding the cellular mechanisms that may be the driving factor for the microstructural development or the breakdown of regional structural integrity. Pathology specific alterations in cerebral metabolite levels have been associated with neurite microstructural alterations [13–15], therefore, it is imperative to investigate the association between POE-mediated cerebral metabolism and microstructural alterations. Such an effort may provide information on cerebral structural development and integrity which may be related to neuronal or axonal health.

In order to elucidate the effects of POE on neurobehavioral outcomes, we developed a mouse model of POE that models a clinical scenario wherein a woman dependent on prescription painkillers (in this case oxycodone) undergoes medication-assisted therapy using methadone and then subsequently becomes pregnant [7]. Clinically, methadone is one of the highest sources of PME [16, 17]. Our prenatal methadone exposure (PME) model recapitulates many of the phenotypes associated with POE. Our previous studies have reported very high methadone levels in fetuses with a precipitous drop in neonates, the expression of neonatal opioid withdrawal, as well as a variety of neurological and behavioral disturbances as a result of PME [7, 9, 18, 19]. We recently performed a neuroimaging assessment of our PME mice where we demonstrated the efficacy of diffusion magnetic resonance imaging (dMRI) in identifying widespread changes in microstructure due to PME [9]. Integrating these microstructural tissue properties with neurometabolites will provide important cell-specific information on the substrate of the cerebral tissue affected by the opioid exposure. We hypothesize that opioid mediated neurodevelopmental changes will influence the regional neurochemistry. We further hypothesize that cerebral metabolite profile will exhibit varying degree of association with regional neurite microstructure in PME offspring. Such a multimodal approach can improve our understanding on the relationship between disturbed cerebral metabolite levels and associated neurite microstructural disorganization. In this study, we are specifically focusing on the dorsal striatum as we have previously identified disrupted neurophysiological processes and biochemical alterations in this brain region [7, 18, 19]. Given the role of dorsal striatum in regulating addiction related behavior and motor control in PME mice [20, 21], it is highly likely that disturbed physiological and neurometabolic changes in this region contribute to the observed behavioral disturbances.

## MATERIAL AND METHODS

### Animal preparation

Our previous protocol was used to produce mice with PME [7]. Briefly, single-housed adult female C57Bl/6J (Jackson Labs, Bar Harbor, ME) mice were treated with either saline (10 ml/kg, s.c. b.i.d.) or an escalating dose treatment with oxycodone (10 mg/kg to 30 mg/kg, s.c.b.i.d.) over the course of 8 days. These oxycodone-treated females were then transferred to methadone treatment (10 mg/kg, s.c.b.i.d.). 5 days after switching to methadone the females were mated with an 8-week-old C57Bl/6J male mouse. These mice continued to receive the same dose of methadone throughout pregnancy and until PME offspring were weaned at postnatal day 28. Saline-treated mice continued to be treated with saline throughout pregnancy to produce prenatal saline exposure (PSE) offspring. Offspring were housed 2-4 mice/cage until neuroimaging was performed at 8-9 weeks of age as previously indicated[9].

All experiments performed with dams, sires, and offspring followed the National Institutes of Health guidelines for the usage of laboratory animals and experimental protocols were approved by the Indiana University School of Medicine Institutional Animal Care and Use Committee. All ARRIVE guidelines were also followed [22].

### MR acquisition

For MR experiments, initially animals were anesthetized under 3% isoflurane in an induction chamber. The anesthetized mice were transferred to an MR compatible cradle and positioned in an MRI compatible head holder to minimize head motion. Anesthesia was then maintained at 1.5% isoflurane in 100% oxygen throughout imaging. Respiration rate was monitored using a pressure pad placed under the animal abdomen and animal body temperature was maintained by a warming pad (37 °C) placed under the animal. The in vivo imaging was conducted on a horizontal bore 9.4 Biospec pre-clinical MRI system (Bruker BioSpin MRI GmbH, Germany) equipped with shielded gradients (maximum gradient strength=660mT/m, rise time = 4750 T/m/s). An 86 mm quadrature volume resonator was used for transmission and a 4-element array cryocoil was used for signal reception (cryoprobe, Bruker, BioSipn). For dMRI, a 2D multi-shell diffusion acquisition scheme was used. The DWIs were acquired using a segmented dual spin-echo echo planar imaging (EPI) sequence using the following parameters: TE/TR = 35.84/3000 ms, δ/Δ = 3/15 ms, matrix size = 128 × 125, voxel size = 130 × 130×130 μm^3^, number of slices = 40, slice thickness = 130 μm, 2 b-value shells (1000 and 2000) s/mm^2^, 10 b0 (5 for each shell), 32 diffusion encoding direction for b = 1000 and 56 for b = 2000 s/mm^2^. A high-resolution FLASH based multi-slice 2D anatomical reference image was acquired to facilitate MRS voxel (1.8 mm^3^) placement in the right dorsal striatum (Figure 1, A and B). The reference scan was acquired using the following acquisition parameters: TE/TR = 3/15ms, slice thickness = 500 μm, in-plane voxel resolution = 78×78 μm^2^, number of slices = 20, number of averages = 3, matrix size = 192×192, flip angle = 10°. To understand the potential contribution of tissue T1 relaxation time on metabolic quantification, a single slice T1 map was acquired. The location of the FoV overlapped with the MRS voxel location. T1 map acquisition is based on single slice Fluid attenuation inversion recovery technique with EPI (FLAIR_EPI), with the following acquisition parameters: TE=17ms, Recovery time = 10000ms, slice thickness = 1000 μm, repetition time (TI) = [30, 100, 200, 300, 400, 500, 600, 700, 800, 900, 1000, 1100, 1400, 1700, 2000, 2300, 2600, 2900, 3200, 3500] ms, matrix size = 128×96 and inversion slab thickness = 4.0 mm.

**Figure 1:**
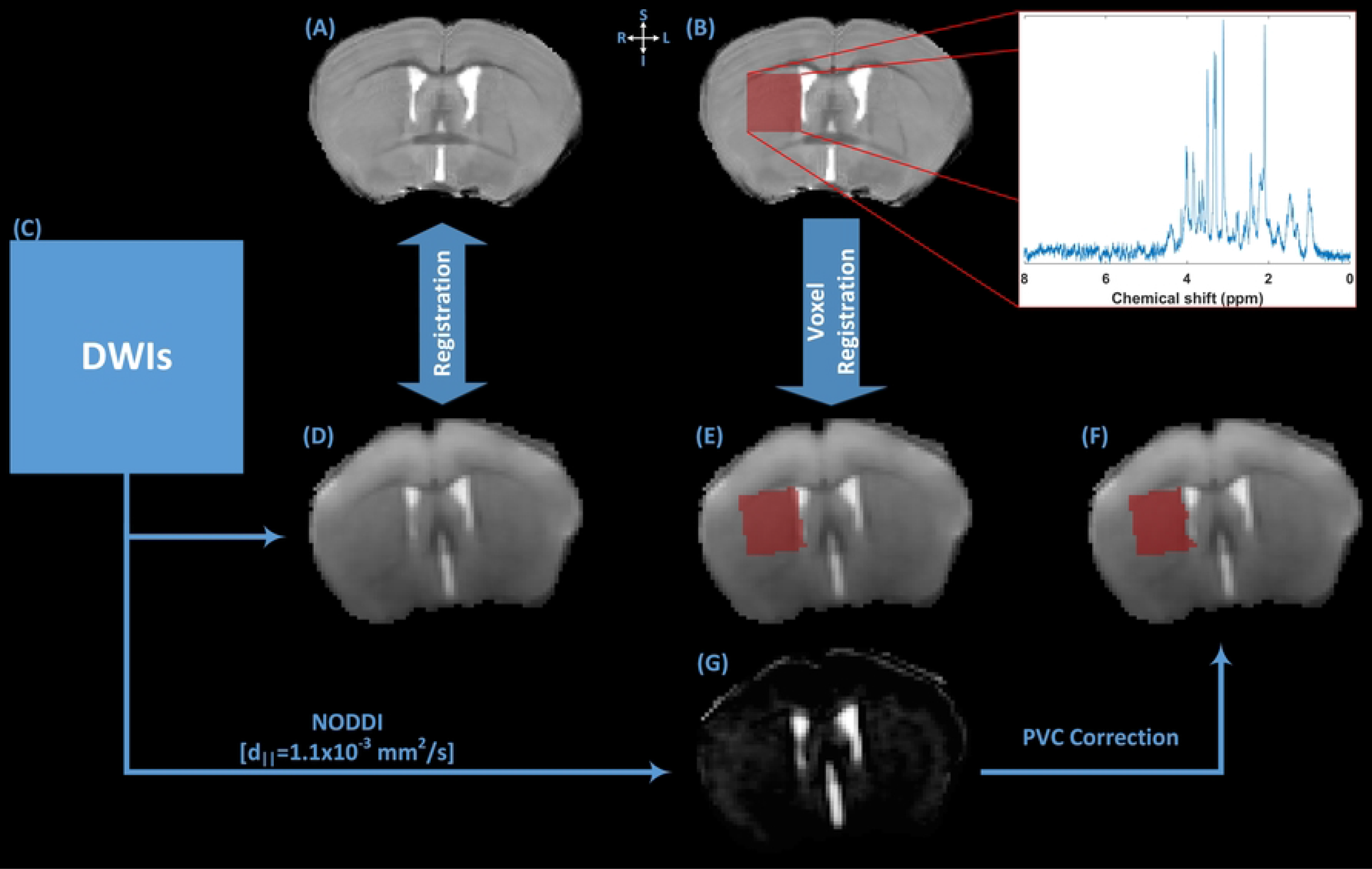
The schematics of the MR image processing framework used in the study. (A) T2-weighted anatomical reference image was used for the manual placement of a 1.8 mm^3^ voxel in the right dorsal striatum region of the animal. (B) T2-weighted anatomical reference image with voxel mask in the right DSR, Inset shows the raw water-suppressed ^1^H-MR spectrum from the single voxel location. (C) DWIs were used to extract averaged b0 image and NOODI derived parametric maps of microstructure. (D) b0 image was linearly register to (A) T2-weighted anatomical reference image. Localized voxel location from (B) T2-weighted anatomical reference image was mapped to subject’s diffusion space using (E) averaged b0 image. (G) Cerebrospinal fluid (CSF) partial volume contamination (PVC) was estimated from NODDI derived VF_CSF_ parameter. (F) CSF PVC was corrected in the mapped voxel. Using the corrected voxel mask, mean values of microstructural parametric maps were calculated for further analysis.

The single voxel spectra (SVS) were acquired with a Stimulated Echo Acquisition Method (STEAM) spectroscopy sequence. A 1.8 mm^3^ voxel was manually placed in the right dorsal striatum using the acquired anatomical reference Scan (Figure 1, B). Before the spectra acquisition, automated localized shimming was performed using Bruker provided FASTMAP method. The field was shimmed to < 15 Hz Full Width Half Maximum (FWHM) linewidth of the unsuppressed water peak. The following parameters were used for spectra acquisition: TE=4ms, mixing time (TM)=10ms, TR=2500 ms, 2048 complex data points, spectral width = 4000Hz, Outer volume suppression (OVS) enabled, and water signal suppression was achieved using variable power and optimized relaxation delays (VAPOR) scheme. OVS slice thickness=5mm, gap to voxel=1 mm, number of dummy scans = 8, number of averages = 384. An additional SVS was acquired as a reference for absolute metabolic quantification using the same acquisition parameters as above except without water suppression enabled. The number of averages for unsuppressed spectra was 10.

### MR data processing

The pre- and post-processing methodologies pertaining to dMRI is described elsewhere [9]. Briefly, the raw DWIs were denoised [23], brain segmented [24], motion and eddy current induced geometric distortion correction [25] and B1 field inhomogeneity corrected [26]. For dMRI model fitting, voxel-based diffusion tensor imaging was performed uing ‘*DTIFIT*’ function in FMRIB Software Library (FSL v6.0) [27] using DWI data from b=0 and 1000 s/mm^2^. Multi-compartment microstructural imaging was performed using Bingham-NODDI model [28]. The high-resolution anatomical reference image was denoised using Non-local Means (*NLMeans*) filter implemented in ANIMA (https://anima.irisa.fr) [29]. Using a custom-built python script, the MRS voxel location was mapped on the reference image (Figure 1, B). To extract microstructural parameters from the MRS voxel location, the denoised reference image was skull stripped [24], bias field corrected [30] and then linearly registered to the b0 image in the diffusion space (Figure1, A, C and D). Using the resulting transformation matrix, MRS voxel mask was translated to b0 image space (Figure 1, B and E). Within the MRS voxel, Cerebrospinal fluid (CSF) partial volume contamination (PVC) was estimated from NODDI derived VF_CSF_ map (Figure 1, G) and image voxels, within MRS voxel, containing PVC were removed (Figure 1, F). This PVC corrected MRS voxel mask was used to extract total MRS voxel volume and mean values of ODI, V_ISO_, VF_IC_ and VF_EC_ using the FSL ‘*fslstats’* tool (Supplementary table 1). T1 relaxation parameters were estimated on Paravision 360 ‘*Image sequence analysis*’ tool using a square region of interest (1.8×1.8 mm^2^) at the approximate location of the MRS voxel with the default settings.

LC Model [31] was used for spectral fitting and absolute metabolite quantification (Figure 2). Before quantification, the raw water suppressed spectra was inspected for poor water suppression, lipid contamination and motion artefacts. The basis-set included model spectra for alanine (Ala), aspartate (Asp), creatine (Cr), phosphocreatine (PCr), γ-aminobutyric acid (GABA), glucose (Glc), glutamine (Gln), glutamate (Glu), glutathione (GSH), glycerophosphpcholine (GPC), phosphocholine (PCh), myo-inositol (mIns), lactate (Lac), N-acetyl aspartate (NAA), N-acetylaspartylglutamate (NAAG), scyllo-inositol (Scyllo) and taurine (Tau). Absolute quantification was performed using the unsuppressed water spectra as a reference. Correction factor related CSF PVC and T1 relaxation of water were also introduced in absolute metabolite quantification. The reliability of the measured metabolite concentration was estimated by Cramer-Rao lower bounds (CRLB) and only those metabolites with CRLB < 15% were selected for further analysis. The metabolite concentrations are reported in μmol/g. PCh and GPC were expressed as total choline (tCh) and Cr and PCr were expressed as total creatine (tCr).

**Figure 2:**
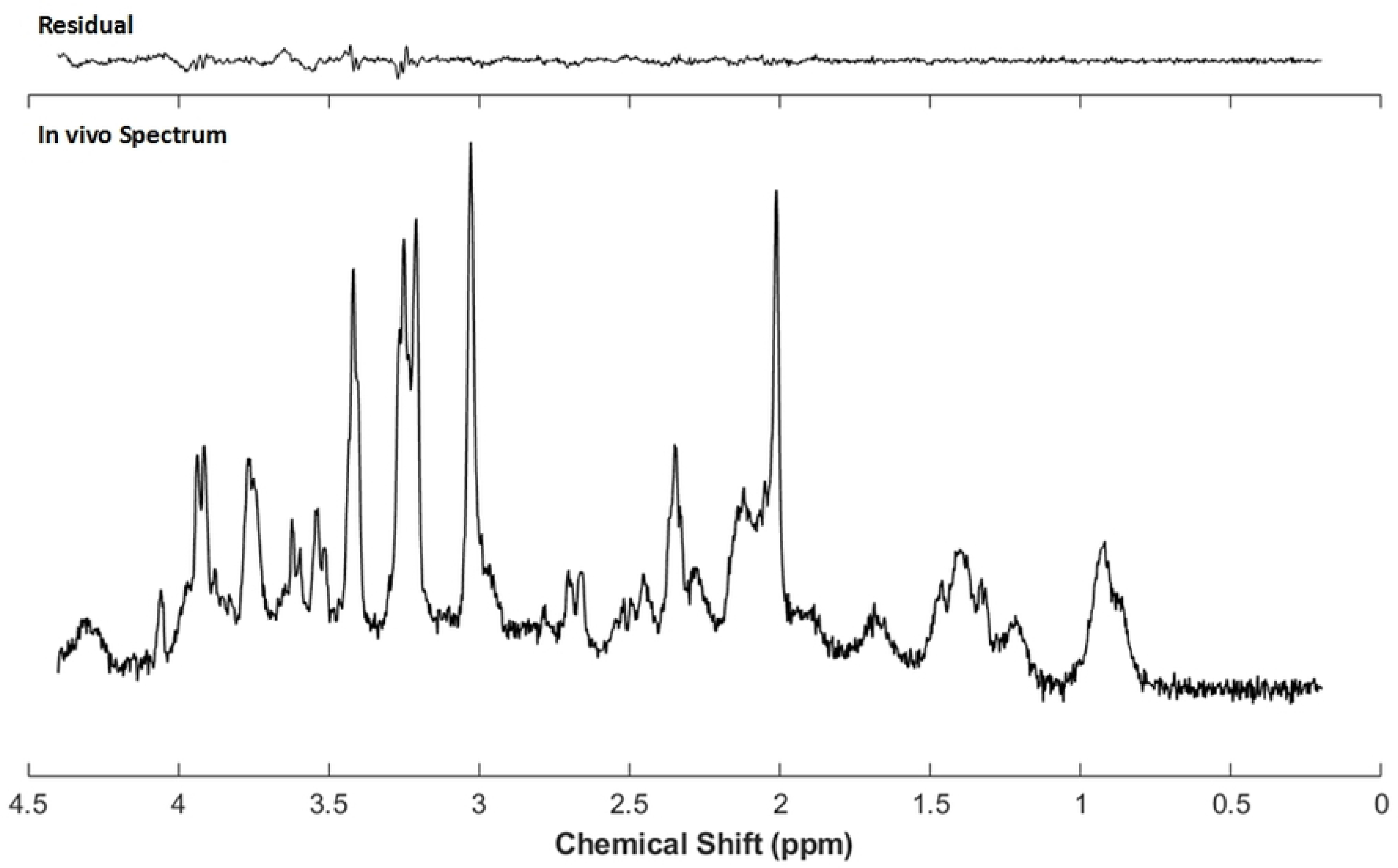
A representative single voxel ^1^H-MR spectrum from a PME mouse using a short echo (TE = 4ms) STEAM sequence at 9.4 T. the In vivo ^1^H single voxel magnetic resonance spectrum was obtained using a 1.8 mm^3^ isotropic voxel located in right dorsal striatum region of the animal. Spectrum quantification was performed using LC Model. Quantification reliability was estimated using Cramer-Rao lower bounds (CRLB). Metabolite data with CRLB > 15% were considered unreliable and exclude from analysis.

### Statistical analysis

To investigate group differences in absolute metabolite concentrations, independent T-test with general linear model (GLM) was used. Benjamini-Hochberg false discovery rate (FDR) correction was used to account for multiple comparison and p_FDR_ < 0.05 was deem significant. To evaluate the relationship between neurochemical levels and microstructural indices in the same region-of-interest (i.e., RDS), partial correlation analyses were performed, and PVC MRS voxel volume was included as a covariate. The analyses were performed in SPSS (IBM SPSS, Version 27).

In order to assess the role of neurometabolites on microstructural organization, we conducted partial correlation analyses between regional metabolite concentration levels and the microstructural parameters in PSE and PME groups, separately. Furthermore, to test, whether the respective correlations in intra-group associations are significantly different from each other, we transformed the individual correlations to Fisher Z scores. Z scores were compared and analyzed for statistical significance by calculating the observed z test statistics (Z_Observed_) [32].

## RESULTS

### Effect of prenatal methadone exposure on cerebral metabolites

Significant group differences between PSE and PME offspring were observed across a number of neurometabolite concentration levels in the RDS (Table 1). PME offspring exhibited significantly lower concentration levels of NAA (p_FDR_ < 0.05; Cohens’d =1.83; 95% CI: 0.53 – 3.08), Tau (p_FDR_ < 0.05; Cohens’d =1.63; 95% CI: 0.37 – 2.83), GSH (p_FDR_ < 0.05; Cohens’d =1.60; 95% CI: 0.35 – 2.81), tCr (p_FDR_ < 0.05; Cohens’d =1.52; 95% CI: 0.29 – 2.70) and Glu (p_FDR_ < 0.05; Cohens’d =1.43; 95% CI: 0.21 – 2.59).

**Table 1.** Comparison of neurometabolite concentrations in right dorsal striatum measured by 1H-MRS.

| Metabolites<br>[μmol/g] | Mean difference<br>PSE-PME | p<sub>FDR</sub> | Effect size<br>(Cohen's d) | 95% Confidence<br>Interval |
| --- | --- | --- | --- | --- |
| NAA | 0.87 | **0.03** | 1.83 | 0.53 – 3.08 |
| Tau | 1.73 | **0.03** | 1.63 | 0.37 – 2.83 |
| GSH | 0.28 | **0.03** | 1.60 | 0.35 – 2.81 |
| tCr | 0.89 | **0.03** | 1.52 | 0.29 – 2.70 |
| Glu | 0.94 | **0.04** | 1.43 | 0.21 – 2.59 |
| Gln | 0.36 | 0.08 | 1.14 | -0.02 – 2.26 |
| GABA | 0.11 | 0.30 | 0.67 | -0.42 – 1.74 |
| Ins | 0.29 | 0.31 | 0.61 | -0.47 – 1.67 |
| tCh | 0.02 | 0.68 | 0.23 | -0.83 – 1.27 |

Abbreviations: PSE = postnatal saline exposure, PME = postnatal methadone exposure, NAA = N-acetylaspartate, Tau = taurine, GSH = glutathione, tCr = total creatine (Cr+PCr), Glu = glutamate, Gln = glutamine, GABA = γ-aminobutyric acid, Ins = myo-inositol, tCh = total choline (GPC + PCh)

### Cerebral metabolite associations with neurite microstructure

Table 2 show the partial correlation between the regional metabolite concentration levels and diffusion microstructural indices in PME group. Total creatine exhibited strong positive associations with regional ODI (r = 0.89, p < 0.05) and with regional VF_IC_ (r = 0.87, p < 0.05). Glu was negatively associated with regional mean diffusivity (r = −0.92, p < 0.01) and was positively associated with regional ODI (r = 0.94, p < 0.01). After FDR correction for multiple comparison, Glu – ODI association remained statistically significant (p_FDR_ < 0.05). No significant correlations were observed between regional neurometabolite levels and regional microstructural parameters in the combined groups (PME+PSE; See Supplementary Table 2) or in the PSE group alone (Supplementary Table 3).

**Table 2:** Partial correlation between the neurometabolite concentrations and the diffusion microstructural metrices in the right dorsal striatum.

| Metabolites<br>[μmol/g] | FA |  | MD (mm<sup>2</sup>/s) |  | ODI |  | VF<sub>IC</sub> |  |
| --- | --- | --- | --- | --- | --- | --- | --- | --- |
|  | r | p value | r | p value | r | p value | r | p value |
| NAA | 0.549 | 0.260 | 0.224 | 0.670 | 0.122 | 0.818 | 0.449 | 0.372 |
| Tau | -0.530 | 0.279 | -0.663 | 0.152 | 0.416 | 0.412 | 0.089 | 0.868 |
| GSH | -0.092 | 0.862 | -0.293 | 0.573 | 0.410 | 0.420 | 0.676 | 0.141 |
| tCr | 0.046 | 0.931 | -0.786 | 0.064 | 0.891 | **0.017<sup>‡</sup>** | 0.873 | **0.023<sup>‡</sup>** |
| Glu | -0.375 | 0.464 | -0.924 | **0.008<sup>‡</sup>** | 0.939 | **0.005<sup>‡</sup>** | 0.741 | 0.092 |
| Gln | 0.741 | 0.092 | 0.351 | 0.495 | -0.090 | 0.865 | 0.403 | 0.428 |
| GABA | 0.427 | 0.398 | 0.043 | 0.935 | -0.084 | 0.875 | 0.273 | 0.601 |
| Ins | 0.062 | 0.908 | -0.698 | 0.123 | 0.749 | 0.087 | 0.652 | 0.160 |
| tCh | 0.317 | 0.540 | -0.649 | 0.163 | 0.715 | 0.110 | 0.686 | 0.132 |

\*Denotes p<sub>FDR</sub> < 0.05; ‡ denotes the respective correlations are significantly different between groups (p<0.01). Abbreviations: r = correlation coefficient, NAA = N-acetylaspartate, Tau = taurine, GSH = glutathione, tCr = total creatine (Cr+PCr), Glu = glutamate, Gln = glutamine, GABA = γ-aminobutyric acid, Ins = myo-inositol, tCh = total choline (GPC + PCh), FA = fractional anisotropy, MD = mean diffusivity, ODI = orientation dispersion index, VF<sub>IC</sub> = intra-cellular volume fraction.

## DISCUSSION

The study employed short echo time STEAM ^1^H-MRS sequence to obtain regional neurochemical profile and used multi-shell dMRI, aided with multi-compartment diffusion modeling, to extract quantitative information related to regional microstructural organization. Using highly resolved short TE spectra, we successfully quantified various intracellular metabolite concentrations, thus were able to probe PME associated biochemical alterations. With these advanced imaging tools, the present study investigated the differences in cerebral metabolite concentration levels in the right dorsal striatum region between PSE and PME offspring at 8 weeks of age. In the PME offspring group, a significant decrease was observed in a number of neurometabolites (including NAA, Taurine, Glutamate, glutathione and creatine) in the right dorsal striatum region compared to the PSE group. We further examined the relationship between regional neurochemistry and microstructural organization in PME offspring and found significant associations between neurometabolite levels and microstructural changes in the PME group. However, we did not find significant associations between neurometabolite concentration levels and microstructural parameters in the control group.

NAA, which is synthesized in mitochondria, is predominantly present in the neuronal component [33]. Higher NAA levels are associated with neuronal health and synaptic integrity [34–37]. Thus, NAA is considered a marker of neuronal integrity and function. The results of our study showed significantly lower levels of NAA in the dorsal striatum of PME group compared to control PSE group. A number of studies in aging [38], Alzheimer’s disease pathology [39] and multiple sclerosis [40] have reported association between neuronal integrity and NAA levels. However, there is a strong argument that at brain maturation and development stage, changes in NAA levels maybe associated with the development of dendrites and synaptic connections [41, 42]. A recent pre-clinical MRS based study suggested that NAA might be a biomarker of neuronal insult after prenatal opioid exposure and subsequent withdrawal [12]. Based on the current evidence, it is possible that lower levels of NAA in PME offspring might be associated with withdrawal. Glutamate is a primary excitatory neurotransmitter in mammalian brain [43]. It is involved in memory and learning, synaptic plasticity, synaptic connections, neuronal growth and differentiation [44]. A number of studies have reported the influence of the dysfunctional glutamatergic system in mood disorders [45] and opioid dependence [46, 47]. Our results showed significant reduction in Glu levels in PME offspring, compared to PSE group. Although the reduction of Glu measured with MRS in PME offspring is from multiple neuronal and glial pools including many metabolic functions in addition to Glu’s role in excitatory neurotransmission, this reduction in Glu could reflect an effect of opioid exposure on Glu synaptogenesis and Glu-mediated neurotransmission. Indeed a recent study by Grecco, Muñoz [18] reported the impaired synaptic Glu signaling pathways in excitatory neurotransmission in the dorsal striatum of adolescent PME offspring. In agreement with healthy aging study [48], our results (supplementary Figure 1) also demonstrated strong positive correlation between Glu and NAA in PSE group, which supports the hypothesis that Glu is an indicator of synaptic health.

Taurine plays an important role in neurodevelopment [38]. The role of Tau in neurite proliferation, synaptogenesis and synaptic transmission is well documented [49]. In our study, Tau concentration levels were significantly lower in PME offspring compared to control group, suggesting long lasting effect of PME on Tau-mediated neuroadaptations related to neurodevelopment [11]. GSH is an antioxidant which is present in brain, primarily in astrocytes [50]. GSH is primarily implicated in oxidation-reduction reactions, serves as a protector against toxic agents such as reactive oxygen species (ROS) [51]. Alterations in cerebral GSH levels from opioid withdrawal mediated oxidative imbalance may reflect inflammatory processes and mitochondrial dysfunction [52, 53]. Our results demonstrated significant reduction in GSH concentration levels on RDS of PME offspring group, compared to the control group. This may suggest the long-lasting effect of methadone on cerebral oxidative imbalance in prenatally exposed offspring.

Total creatine plays a very significant role in cellular energy metabolism. In central nervous system, Cr and PCr are present in neurons and glial cells [54]. Alterations in cerebral tCr levels indicate mitochondrial dysfunction [55] and disturbed cerebral energy metabolism [56]. To date, there are conflicting reports on altered bioenergetics in human opioid and methadone maintenance therapy studies [46]. Our results indicate significant reduction of tCr levels in PME offspring, compared to control group. These results may suggest long lasting effects of prenatal opioid exposure on cerebral bioenergetics. Furthermore, our results indicate that tCr should not be considered as an internal reference in PME studies due to significant group-wise alterations in its concentration levels. Moreover, the metabolite ratio is less sensitive when the concentration of both metabolites is affected by considered pathology.

Previous studies in healthy aging suggest the association between regional neurochemical composition and microstructural organization [57, 58]. In this study we observed a strong positive association between regional Glu levels and ODI in PME group. Interestingly, the trajectory of this association in PSE group was in opposite direction and was not statistically significant. Total Cr in PME group also exhibited positive associations with regional ODI and VF_IC_ parameters, although these correlations did not survive the FDR based corrections but do suggest the possible involvement of altered neurochemical composition in regulating neurite microstructural organization in PME offspring.

There are some limitations which need to be highlighted. Given the exploratory nature of the work, we only scanned male offspring. We also only scanned the right dorsal striatum, and other brain regions certainly warrant further investigation. In addition, the sample size was relatively small and therefore, generalization of these results should be considered with caution. Additionally due to the cross-sectional nature of the study design, it was not possible to establish cause and effect relationship between cerebral metabolites and microstructural parameters. Despite these limitations, the study suggests that PME can significantly affect neurodevelopment in PME offspring by perturbing neurometabolism and brain microstructure and the effects of these perturbation can persist into late adolescent and early adulthood.

## ACKNOWLEDGEMENTS

This work was supported in part by the Stark Neurosciences Research Institute, Indiana University (BKA) and NIH/NIAAA F30 AA028687 (GGG).

## DATA AVAILABILITY STATEMENT

The datasets used and/or analyzed during the current study are available from the corresponding author on reasonable request.

